# Lumbar endplate microfracture injury induces Modic-like changes, intervertebral disc degeneration and spinal cord sensitization – An In Vivo Rat Model

**DOI:** 10.1101/2023.01.27.525924

**Authors:** Dalin Wang, Alon Lai, Jennifer Gansau, Alan C. Seifert, Jazz Munitz, Kashaf Zaheer, Neharika Bhadouria, Yunsoo Lee, Philip Nasser, Damien M. Laudier, Nilsson Holguin, Andrew C. Hecht, James C. Iatridis

## Abstract

**BACKGROUND CONTEXT**: Endplate (EP) injury plays critical roles in painful IVD degeneration since Modic changes (MCs) are highly associated with pain. Models of EP microfracture that progress to painful conditions are needed to better understand pathophysiological mechanisms and screen therapeutics.

**PURPOSE**: Establish in vivo rat lumbar EP microfracture model with painful phenotype.

**STUDY DESIGN/SETTING**: In vivo rat study to characterize EP-injury model with characterization of IVD degeneration, vertebral bone marrow remodeling, spinal cord sensitization, and pain-related behaviors.

**METHODS**: EP-driven degeneration was induced in 5-month-old male Sprague-Dawley rats L4-5 and L5-6 IVDs through the proximal vertebral body injury with intradiscal injections of TNFα (n=7) or PBS (n=6), compared to Sham (surgery without EP-injury, n=6). The EP-driven model was assessed for IVD height, histological degeneration, pain-like behaviors (hindpaw von Frey and forepaw grip test), lumbar spine MRI and μCT analyses, and spinal cord substance P (SubP).

**RESULTS**: EP injuries induced IVD degeneration with decreased IVD height and MRI T2 values. EP injury with PBS and TNFα both showed MC type1-like changes on T1 and T2-weighted MRI, trabecular bone remodeling on μCT, and damage in cartilage EP adjacent to the injury. EP injuries caused significantly decreased paw withdrawal threshold and reduced grip forces, suggesting increased pain sensitivity and axial spinal discomfort. Spinal cord dorsal horn SubP was significantly increased, indicating spinal cord sensitization.

**CONCLUSIONS**: EP microfracture can induce crosstalk between vertebral bone marrow, IVD and spinal cord with chronic pain-like conditions.

**CLINICAL SIGNIFICANCE**: This rat EP microfracture model of IVD degeneration was validated to induce MC-like changes and pain-like behaviors that we hope will be useful to screen therapies and improve treatment for EP-drive pain.

## Introduction

Chronic back pain is a prevalent musculoskeletal disorder and a major cause of disability with enormous socioeconomic burdens worldwide [1–4]. Back pain is highly associated with endplate (EP) defects in relation to Modic changes (MCs) and intervertebral disc (IVD) degeneration PMID: [5–7]. MCs are magnetic resonance imaging (MRI) evidence of inflammatory and fibrotic vertebral bone marrow lesions that associate with adjacent IVD degeneration and EP defects [5, 8–11]. EP defects can be the result of peripheral avulsive fracture or central accumulating microfractures at the vertebral EP [12]. Many clinical studies identify strong associations between EP defect changes as seen on spinal imaging and pain presence, indicating the importance of these clinically-relevant spinal changes [8, 9, 11, 13]. Pain may result from EP microfracture when it accumulates into larger EP defects with bone marrow involvement characterized as MCs; MCs result in the crosstalk between multiple spinal tissues with an autoimmune response of the bone marrow against IVD and nervous system tissues [1, 14–20].

EP-driven and annulus fibrosus (AF)-driven IVD degeneration are often mixed and interacting in clinical back pain patients, so animal models are required to identify the causes and to screen potential treatments for these pain sources that can be distinct [1, 21, 22]. Vertebral EPs and outer annulus fibrosus (AF) are innervated with an abundance of nociceptive neurons that connect to dorsal root ganglion (DRG) and spinal cord. EP defects (or microfracture injury) and elevated pro-inflammatory cytokines from IVD degeneration may irritate nerves and induce pain. Such crosstalk between spinal tissues is particularly important since DRG and spinal cord changes can occur from IVD injury and degeneration without obvious vertebral involvement [23–28]. The clinical importance of MCs motivates a strong need to develop animal models of EP injury. An EP injury (or EP-driven IVD degeneration) model is required to determine if EP microfracture is sufficient to trigger MCs, IVD degeneration and pain, and to study the pathophysiology of MC etiology [17, 19, 29, 30]. Furthermore, an animal model of EP injury is needed to identify therapeutic targets and develop improved treatment strategies for chronic EP-driven back pain.

Animal models for studying EP injuries have been limited to large animals (i.e. porcine and ovine), with outcome measurements restricted to IVD degeneration, and the effects of EP defects on MCs [31–33]. Large animal models involve a high cost of breeding and housing, and assays to characterize pain-related behaviors are not well-established. Rat models are commonly used to study painful spine conditions because they are relatively inexpensive, have fast healing times, exhibit anatomical and biomechanical similarities to the human spine, and have well-characterized behavioral assays to characterize pain-like conditions [24, 28, 34–42]. Rat models are also sufficiently large to enable spine surgical procedures to be accurately performed. However, even in large animal models [31–33], EP defect injuries caused variations in IVD degeneration severity, highlighting a need for precisely performed EP injury creation in any animal model.

The objectives of this study were to i) establish and characterize a lumbar EP microfracture injury model in rats in vivo, ii) evaluate pain-related behaviors and the microstructural characteristics of IVD degeneration and bone marrow remodeling following EP microfracture injury, iii) investigate the changes in spinal cord following EP microfracture, and iv) establish associations for EP injury between MRI NP T2 relation time with IVD degeneration, spinal cord sensitization and pain-related behaviors. Rat lumbar IVDs and vertebral body were evaluated using post-mortem MRI and μCT from sham to injury status. This study provides insight into how EP microfracture injury can progress to MCs, IVD degeneration, spinal cord sensitization and pain-like behaviors, and provides an animal model that induces MC-like changes in order to provide a screening tool for potential therapeutic interventions.

## Materials and methods

### Study design

All experimental procedures were approved by the Institutional Animal Care and Use Committee at the Icahn School of Medicine at Mount Sinai. Nineteen 5-month old male Sprague-Dawley rats (Charles River Laboratory, Wilmington, MA) were randomly divided into 3 groups: Sham (n=6), EP+PBS (n=6), EP+TNFα (n=7). EP+PBS and EP+TNFα groups had EP microfracture injury followed by an intradiscal injection of PBS and TNFα, respectively (Figure 1). For the sham group, vertebral bodies between L4-L6 as well as IVD levels L4-5 and L5-6 were exposed without any injury. Animals were evaluated for pain-related behaviors throughout the 8 week experimental duration (Figure 2), and were otherwise allowed unrestricted movement in cages. The lumbar spine and spinal cord were then assessed using faxitron for IVD height, histology and IVD degeneration scoring, MRI, μCT, and spinal cord sensitization.

**Figure 1:**
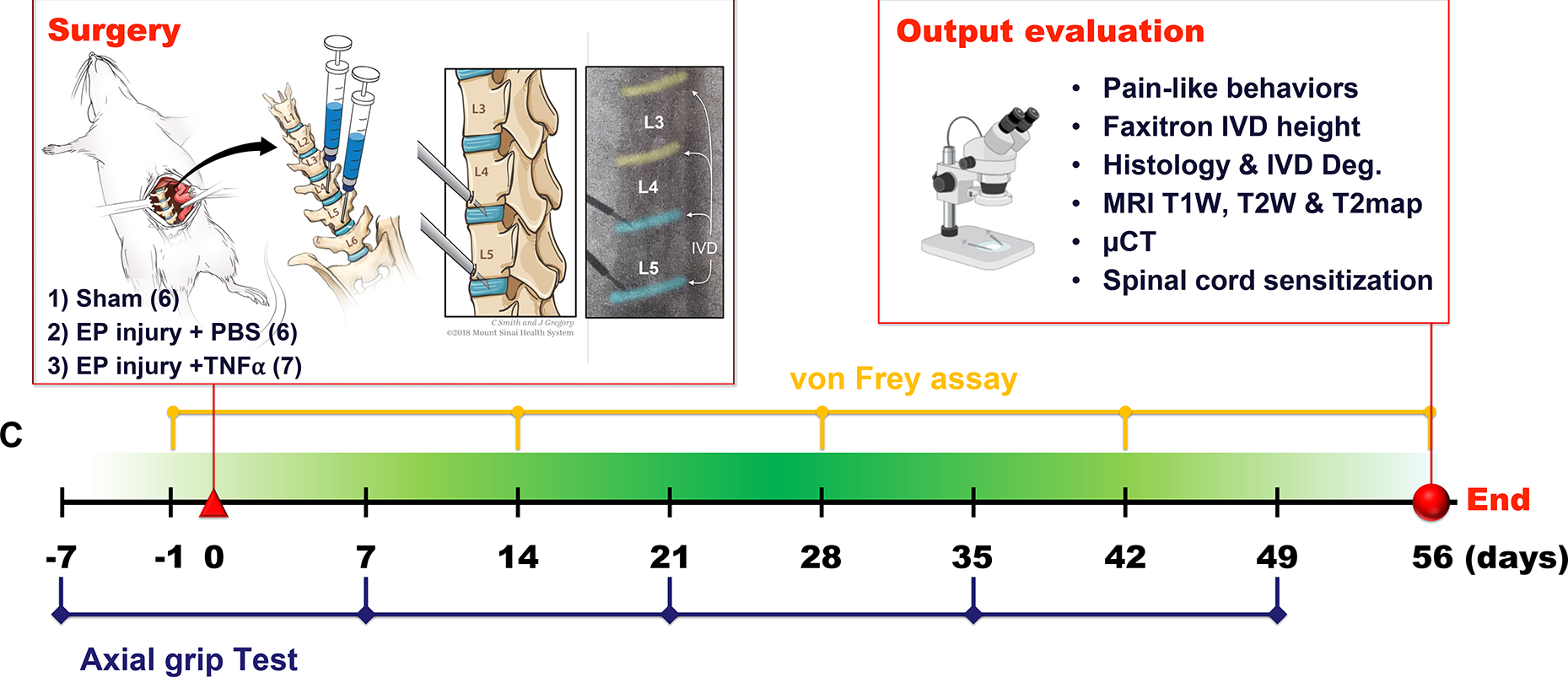
EP microfracture in-vivo model and study design. A) Schematic of procedure with anterior approach. Experimental groups included Sham (n=6); EP injury + PBS injection (n=6) and EP injury + TNF⍺ injection (n=7). B) Output variables after t=56 days (8 weeks). C) Timeline of behavioral measurements.

**Figure 2:**
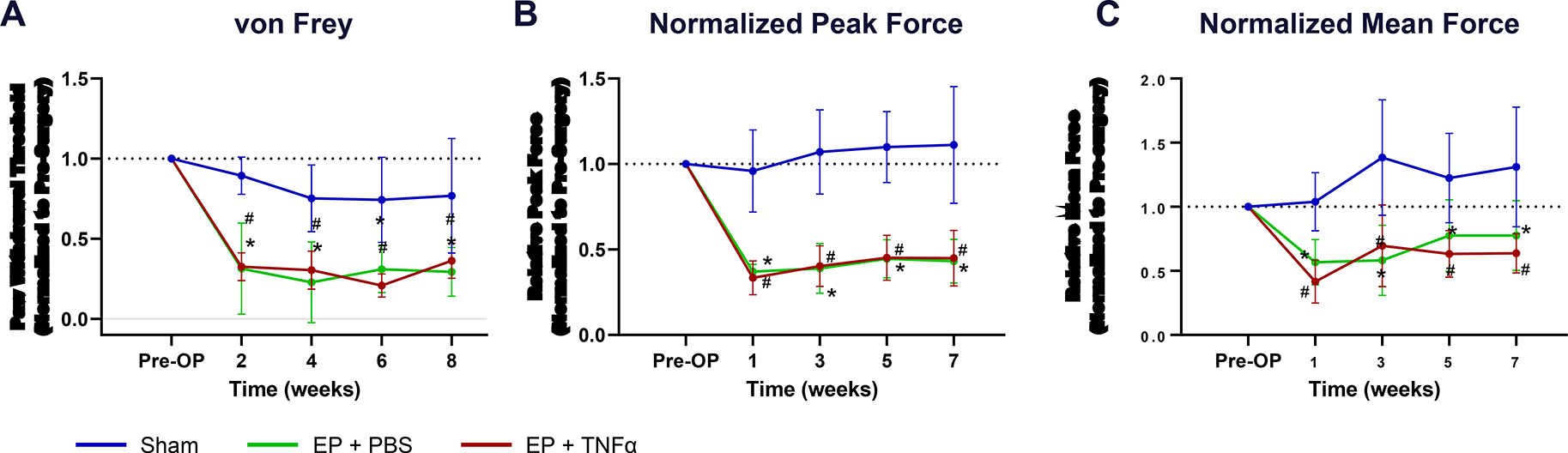
Behavioral testing demonstrates axial sensitivity and central sensitization. A) Paw withdrawal threshold after von Frey test; B) Normalized Peak force and C) normalized mean force after grip test over time # EP+PBS compared to Sham with p < 0.05, * EP+TNFα compared to Sham with p < 0.05.

### Surgical procedure and EP microfracture injury

Surgical procedures were performed under aseptic conditions and general anesthesia via 2% isoflurane (Baxter, Deerfield, IL) [24]. An anterior abdominal incision was used to expose L4~L6 lumbar spine. IVD level was preliminarily identified using preoperative anterior-posterior X-ray images, and confirmed by intraoperative C-arm. Rats underwent either a sham surgical procedure or EP puncture surgery of the L4-5 and L5-6 IVDs. For the EP injury groups, the proximal EPs of L4-5 and L5-6 IVDs were punctured obliquely from the vertebral body at 1.5 mm proximal to the edge of IVDs using a 0.6 mm K-wire, which was controlled by a 3 mm depth stopper (Figure 1). All intradiscal injections were performed following the EP injury using a 26-gauge needle with a 3 mm depth stopper. A total of 2.5 ul of PBS or TNFα (0.25 ng in 2.5 ul) (80045RNAE50; Sino Biological Inc., Beijing, China) [24, 43] was then slowly injected into each IVD using a calibrated microliter syringe (Hamilton Company, Reno, NV, USA) following the EP injury. All EP injuries were guided and confirmed radiologically using the C-arm (Figure 1C).

Animals were then housed 2 per cage and maintained at a 12/12 hour light/dark cycle (light stage: 7 am to 7 pm) for the experimental duration. Animals were allowed unrestricted movement in cages for the entire experimental duration and co-housed two per cage with the exception of the 24 h post-operative period, when animals were singly housed [28].

### von Frey and axial grip behavioral testing

Pain-related behaviors were evaluated using von Frey assay for hindpaw mechanical allodynia at 0, 2, 4, 6 and 8 weeks post-injury, and grip test for axial lumbar discomfort at 0, 1, 3, 5 and 7 weeks post-injury (Figure 2). The behavioral tests were performed by a single experimenter in a dedicated behavioral analysis room with regular indoor lighting.

The mechanical allodynia at hindpaws was assessed using von Frey assay [24, 25, 44];[40]. All rats were acclimated to handling and test cages for 7 consecutive days before testing. On the day of testing, the rats were acclimated in the test cages for 20 min before testing. Von Frey filaments ranging in force between 0.4 and 26.0 g were applied to the plantar surface of each hindpaw in ascending force, with each filament applied five times. The lowest force filament eliciting nocifensive behaviors in 3 out of 5 applications was identified as paw withdrawal threshold. Nocifensive behaviors included paw licking, extended paw withdrawal, and fanning/ shaking of the paw. The paw withdrawal thresholds from the left and right hindpaws were averaged for statistical analysis.

Axial lumbar discomfort was assessed using a grip strength test on the forepaws as described by literature [45]. Grip strength was measured using a custom-built testing apparatus with a stainless steel grid connected to a uniaxial force sensor. During testing, the animal was gently positioned and allowed to grab the metal grid with both forepaws. The tail of the animal was held and gently pulled until the animal released the grid. This action stretches the lumbar spine, and was therefore considered a measure of axial discomfort. The force data was sampled and recorded for 30 seconds using LabVIEW (National Instruments), and the peak force and mean force were calculated via the analysis of the recorded loading curves. This grip force test procedure was repeated three times and each trial was followed by 10 minutes resting with the rats in their own cages. Results were averaged from the three trials at each time point for statistical analysis.

### Faxitron analysis of disc height and specimen collection

Changes of IVD height were quantified in vivo using faxitron radiography pre-operatively, and at 8 weeks after injury with the animal anesthetized and carefully placed on its side (Figure 3A). At 8 weeks post-surgery, all rats were transcardially perfused with 10% buffered formalin phosphate (Fisher Company, Fair Lawn, NJ, USA) under the condition of anesthetization, both lumbar spinal cord and lumbar spines were dissected and fixed in 10% buffered formalin phosphate. The formalin-fixed spinal cord was used for immunohistochemical analysis for substance P; while the fixed spine was for post-mortem MRI and μCT, followed by histological analysis.

**Figure 3:**
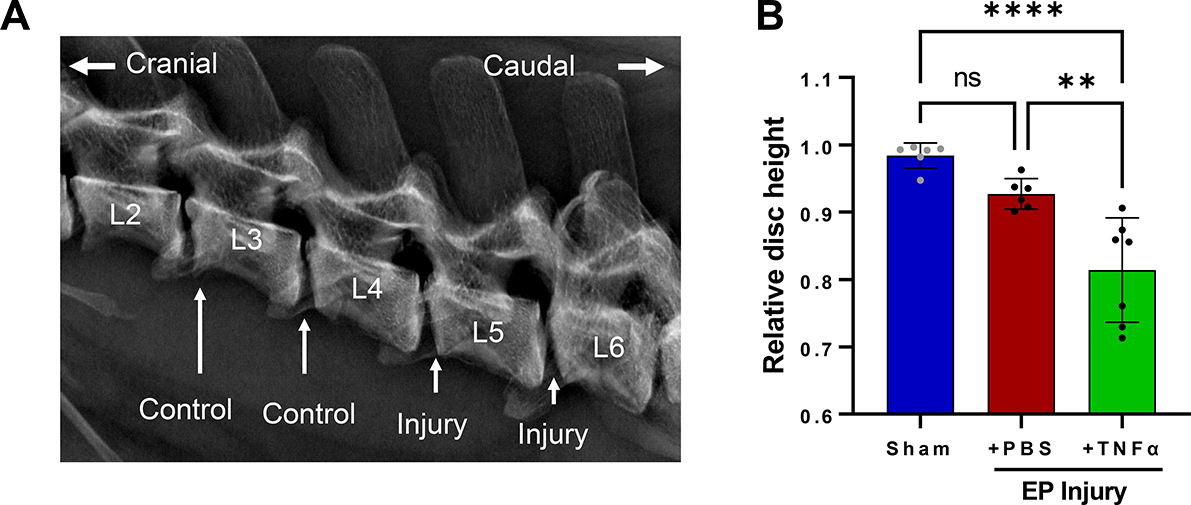
EP Injury causes IVD degeneration. A) Faxitron imaging and B) IVD height loss with injury. **and **** indicate significant differences with p<0.01 and p<0.0001 respectively.

### MRI scan and analysis

MRI was performed on a 9.4T vertical-bore micro-MRI system (Bruker Avance III 400) using a 20-mm quadrature birdcage RF coil (Rapid Biomedical). T1-weighted (T1w) images were acquired using 3D MP-RAGE (139 μm isotropic resolution, TR=4s, TI=1.1s), T2-weighted (T2w) images were acquired using 3D RARE (139 μm isotropic resolution, TR=2s, TE=17ms), and T2 mapping data were acquired using multi-echo 3D spin-echo (remmiRARE, 250 μm isotropic resolution, TR=520ms, TE1=6ms, ΔTE=5ms, 32 echoes). T1w and T2w images were analyzed for Modic changes using methods previously described [10, 46].

T2 maps were produced by fitting a single exponential decay to each voxel’s echo train. Using co-registered anatomical images as a guide, 3D regions of interest (ROI) in the NP, whole IVD, and bone marrow were defined on T2 maps as binary masks using FSLEyes (https://fsl.fmrib.ox.ac.uk/fsl/fslwiki/FSLeyes). Mean values for T2 relaxation time were quantified within each mask, excluding voxels where quality of fit was poor due to insufficient signal (e.g., bone), using fslstats, a utility contained within FSL [47, 48]. The 3D ROIs for NP and whole IVD T2map masks were manually drawn in FSLEyes to enclose the entire NP and IVD using T2w and T1w co-registered images that easily identified the borders of these anatomic structures. The 3D ROI for the bone marrow was created using a 5-voxel cube centered on the endplate defect region, which was then manually cropped to exclude non-marrow tissues (e.g., IVD and cortical bone). The 3D ROIs for the bone marrow High Intensity Zone (HIZ) was created using a 3 voxel cube centered on the brightest area of the EP defect, and on comparable region in the sham animals (which did not exhibit defects), and these smaller ROIs did not require cropping to exclude non-marrow tissues.

### μCT scan and analysis

Vertebral body remodeling was assessed via μCT at 8 weeks post-surgery. μCT was performed on a Nanoscan PET/CT system (Mediso Co) at energy 84 uAs, slice thickness of 0.02 mm, isotropic voxel size at 0.25 mm. Image analysis was performed using Osirix MD, version 11.0. After opening reconstructed CT images, scans were viewed in a sagittal orientation, and 3 volumetric ROIs were hand drawn in each vertebra. The Injury site ROIs characterized the endplate regions, and were drawn using the pencil tool extending from the inner corona of the injury area extending 1 mm along the endplate, and extended horizontally alongside the trabecular area of the bone. For Adjacent ROI regions, sections were drawn beginning 1 mm above the Injury site, and extended 1 mm in height, and horizontally along the width of the trabecular region. The Far Field ROIs were drawn mirroring the size and position of the endplate region, on the endplate on the opposite side of the vertebral body. The 3D ROIs were modeled, and the volume was noted. A histogram of pixel-binned values was created, and all values above 1000 Hounsfield counts (experimentally determined to represent trabecular bone) were counted as bone volume (BV) voxels, and this was compared to the total voxel counts within the ROI, or total volume (TV). BV/TV was calculated, and presented in comparison between the three regions selected for analysis.

### Histology, IVD degeneration score, and immunohistochemical analyses of spinal cord

After the MRI and μCT scanning, the fixed specimens were decalcified, embedded in resin, and sectioned sagittally at 5 μm intervals. The midsagittal section with the EP microfracture were identified, and stained with Safranin-O/fastgreen/hematoxylin for disc morphology and glycosaminoglycan (GAG) content; and with hematoxylin and eosin for IVD cellularity. Slides were then imaged using bright-field microscopy (Leica Microsystems, Inc, Deerfield, IL, USA). IVD degeneration score was determined using a grading system that evaluated NP morphology, NP cellularity, NP-AF border, AF morphology, and EP irregularity [49]. IVD degeneration scoring was performed with three evaluators, who were blinded to the experimental groups, and the degeneration score from the three evaluators were averaged for statistical analysis.

The formalin-fixed spinal cord were paraffin-embedded and sectioned sagittally at 5 μm intervals. Two sections per animal spread across the lumbar spinal cord were selected. After deparaffinization and rehydration, the spinal cord sections were treated with antigen-retrieval buffer, Histo/zyme (H3292, Sigma-Aldrich, Inc, St. Louis, MO, USA), and protein blocking buffer, 2.5% normal horse serum (S-2012, Vector Laboratories, Inc, Burlingame, CA, USA). Sections were incubated at room temperature for 1 hour with mouse monoclonal primary antibodies against rat substance P (1:300 dilution, ab14184, Abcam, Cambridge, MA, USA) or normal mouse serum (ab7486, Abcam) as negative control [24]. After incubation with RTU biotinylated goat anti-mouse IgG secondary antibody (BP-9200, Vector Laboratories, Inc), the sections were treated with DyLight 488 horse anti-goat IgG antibody (DI-3088, Vector Laboratories, Inc). The sections were then Nissl-stained to visualize the neurons and glia cells, washed and mounted (ProLong™ Gold Antifade Mountant with DAPI, P36931, Thermo Fisher Scientific, Waltham, MA, USA). Images were taken at 20x magnification using Leica DM6 B microscope (Leica Microsystems, Inc). Identical microscope settings were used throughout. All images were analyzed using ImageJ, an immunoreactivity (ir) threshold was set and the percentage of SubP-ir relative to area of spinal dorsal horn was quantified, and then averaged between left and right dorsal horn as well as across the two sections from each animal.

### Statistical analysis

All post-injury data of IVD heights, paw withdrawal thresholds and grip strengths were normalized to pre-injury values and presented as percent change to minimize individual variability. Normalized IVD height, spine MRI, μCT, IVD degeneration score, and spinal cord immunohistochemistry were analyzed using one-way ANOVA with Tukey’s multiple comparison test. Pearson’s correlation analyses identified associations between mean NP T2 relaxation times with IVD degeneration grade, spinal dorsal horn SubP-ir, paw withdrawal threshold, and grip force All statistical analyses were performed using Prism (GraphPad, LaJolla, CA), with significance as p < 0.05.

## Results

### Surgery did not affect rat general health

Both sham surgery and EP injury procedures were well-tolerated by the rats. The rat body weight averaged 561±34g, 568±33g, 579±29g, 592±33g, and 611±35g, for pre-surgery and post-surgery weeks 2, 4, 6, and 8, respectively. There were no significant differences between groups at each time point. No obvious stress or discomfort were observed from the general physical examination.

### EP microfracture induced back pain-related behavior

Both groups involving EP injury with either PBS or TNFα had significantly decreased paw withdrawal threshold at hindpaw compared to Sham (Figure 2A), demonstrating increased mechanical sensitivity at the hindpaw and suggesting central sensitization. Both injured groups had significantly decreased forelimb peak grip force and mean grip force compared to Sham at all postoperative timepoints (Figure 2B and C), demonstrating increased axial discomfort. Sham rats did not show significant changes in pain-related behaviors with time. Results of the von Frey and axial grip test together suggest increased pain sensitivity after both EP injury types.

### EP microfracture induced IVD degeneration

At 8 weeks post-surgery, changes in IVD height were significantly different among the three groups: At L4-5, EP+TNFα group decreased to 86.3%±4.9% of baseline, EP+PBS group decreased to 92.8%±3.4%, while the Sham group increased slightly to 100.4%±2.4%. At L5-6, EP+TNFα group decreased to 76.5%±11.5% of baseline, EP+PBS group decreased to 92.5%±3.1%, while the Sham group decreased slightly to 96.7%±2.7%. There was no apparent change in IVD height in the Sham group or at internal control levels (ie, L2-3 and L3-4) that were not injured (Supplementary Figure 1).

Normal IVD morphology was observed in sham surgery animals, while EP injuries with intradiscal injections of PBS or TNFα induced moderate to severe IVD degenerative changes, including smaller and more fibrous NP, decreased number of NP cells, less distinct NP-AF boundaries, disorganized AF lamellae, and observable EP disruptions (Figure 4A). Some NP of EP-injured IVDs were herniated into the adjacent vertebra through the puncture track. There were minimal to no AF tears or disruptions as the AF was kept intact during IVD injury. The semi-quantitative degeneration grading system showed that Sham IVDs had low Total IVD degeneration scores (2.0±2.5, Figure 4B). IVD degeneration scores of both EP+PBS (9.4±1.2) and EP+TNFα (11.6±1.6) injury groups were significantly higher than that of the Sham group (p<0.05). The scores of subcategories of NP morphology, NP cellularity, NP-AF border and EP of the EP injury groups were significantly higher than the Sham group.

**Figure 4:**
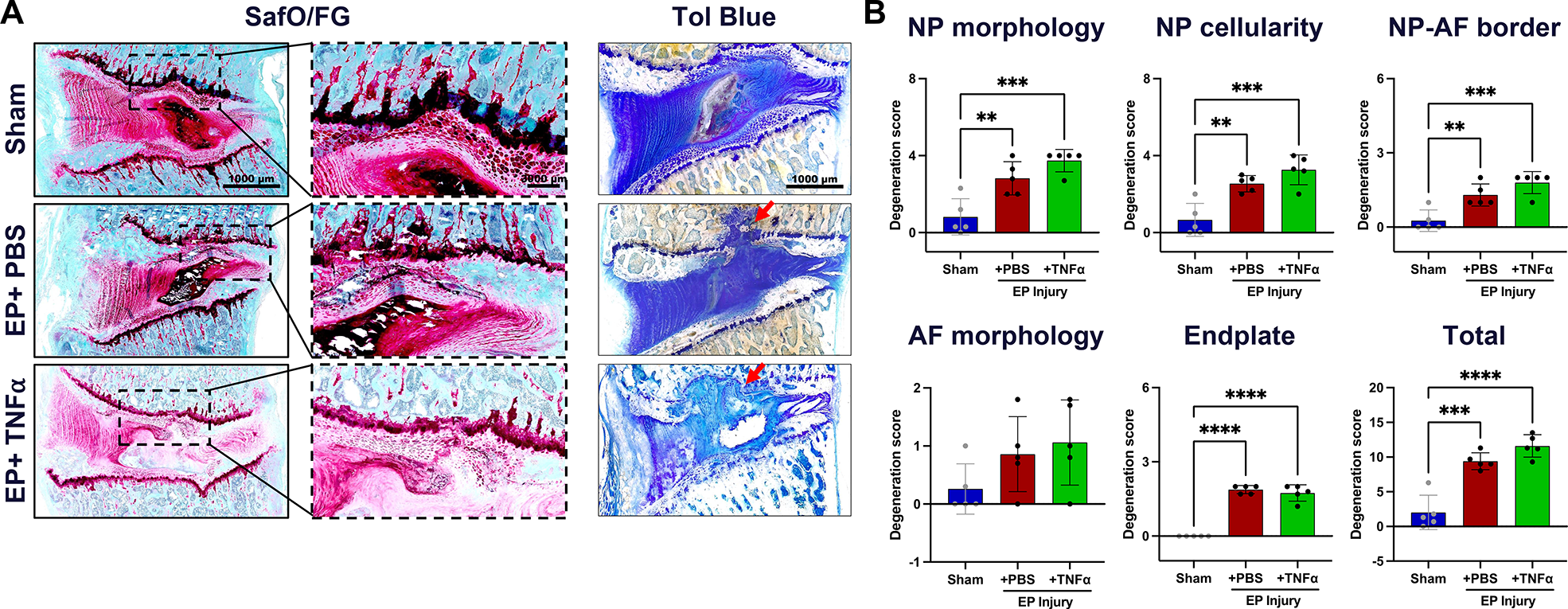
Histology with thin sections stained with SafO/FG and thick ground and polished sections stained with Tol Blue. Red arrows indicating EP defect. B) IVD degeneration grading using Scoring System [49]. **, *** and **** indicate significant differences with p<0.01, p<0.001 and p<0.0001 respectively.

MRI measures of IVD degeneration were performed with mean T2 relaxation time, calculated from T2maps. NP T2 relaxation times tissue significantly decreased with EP+TNFα compared to Sham and EP+PBS, respectively (Figure 5). T2 values of the whole IVD were not affected by the injury, highlighting the localized nature of this injury (Figure 5).

**Figure 5:**
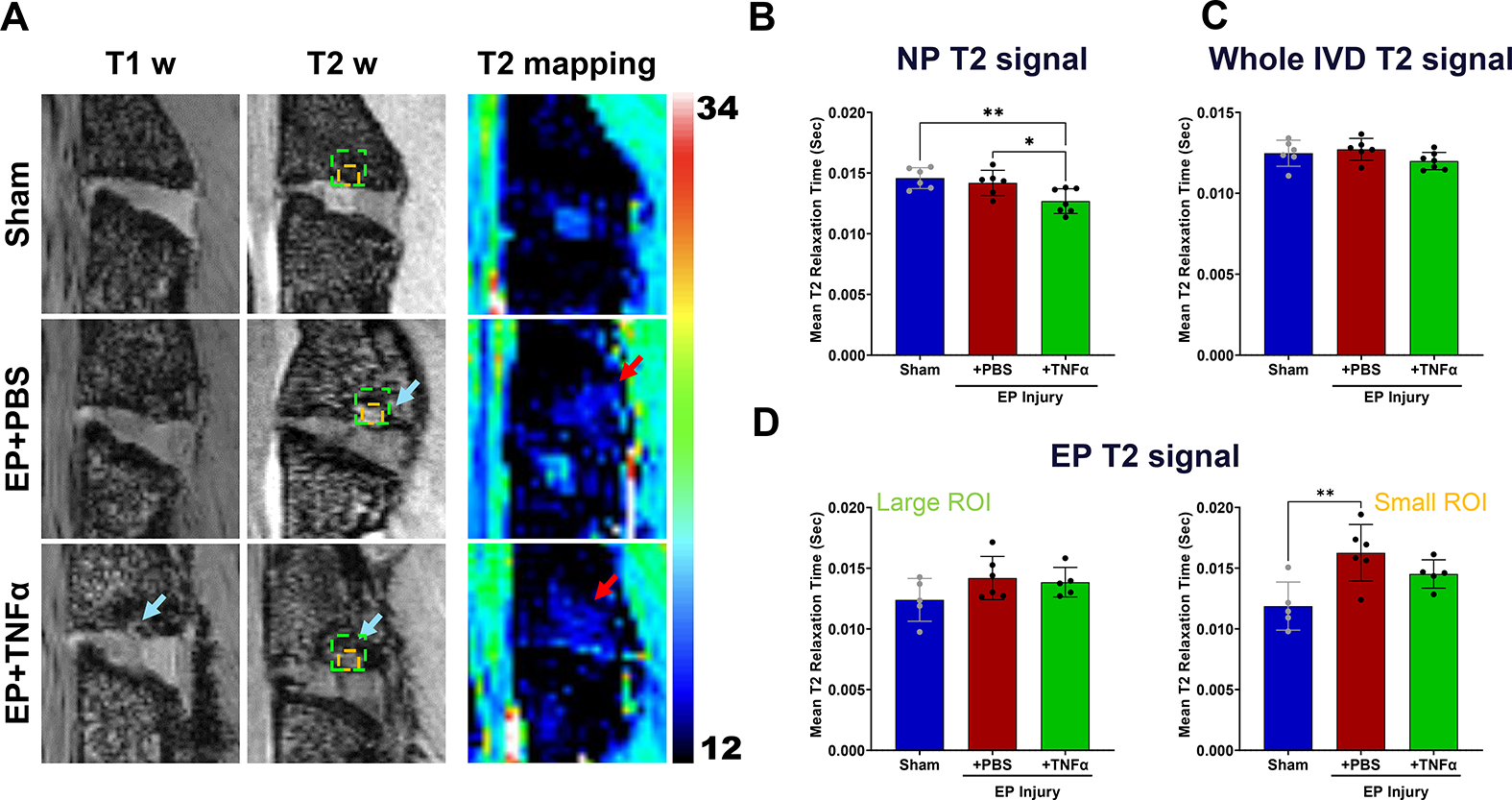
MRI analyses show significant injury following EP injury for both PBS and TNF α. A) Hypointensity on T1w and T2w images (blue arrows) and increased T2 relaxation time (red arrows) are visible around EP defects. B) Mean T2 relaxation time of NP decreased with injury. C) There was no difference in mean T2 relaxation time of the whole IVD among the 3 groups highlighting that this is a localized injury. D) Mean T2 relaxation time at the EP looking at a larger area (left) and more focused HIZ area (right) in which a significant difference between sham and EP=PBS was found. * and ** indicate significant differences between groups with p<0.05 and p<0.01, respectively.

### EP microfracture induced Modic-like changes and bone remodeling

Post-mortem MRI at 8 weeks showed IVD degeneration and bone marrow signal changes. Modic-like changes were visible in both EP injury groups with hypointensity on T1w images and hyperintensity on T2w images in the both injury groups (Figure 5A). Interestingly, T2 mapping sequences indicated reduced T2 relaxation time in the NP region of the EP+TNFα group but not for the whole IVD indicating reduced NP water content and greatest severity of IVDD (Figure 5B and C). Despite the obvious anatomical changes observed on T1w and T2w imaging with the EP defect, the T2 mapping showed little, if any differences in T2 relaxation times. A small ROI focused on the EP HIZ detected a significantly increased T2 relaxation time for the EP+PBS group indicating increased water content suggesting inflammation. The Modic-like changes observed in the T1w and T2w imaging on the EP+TNFɑ group showed larger and more diffuse areas of EP remodeling and inflammation, which are somewhat consistent with the slightly lower values of T2 relaxation time in the HIZ.

The μCT analyses showed trabecular bone remodeling in both EP injury groups (Figure 6A and B), and cartilage EP secondary damage in the EP+TNFα group that was most obvious on histological images (Figure 4A). BV/TV (%) decreased after EP injury compared to Sham at the injury site, adjacent site, and far field in the vertebrae regions (Figure 6C). No significant differences but trends were detected at the injury and adjacent site (p<0.1), likely due to the limited sample size on this analysis. On the far field site, however, a significantly lower BV/TV was found between sham and EP+TNFα (p<0.05).

**Figure 6:**
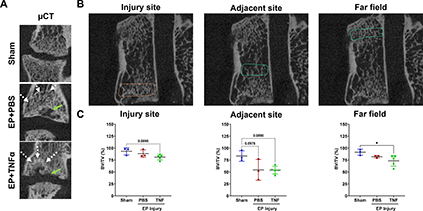
μCT analyses show vertebral disruption and remodeling at the injury site that is not impacted adjacent or at the far field. A) EP defects are highly visible on μCT, and mid-coronal plane of injured lumbar spinal regions vertebrae show trabecular bone remodeling (white arrow) around EP defect (green arrow). B) Quantitative μCT analyses show extensive injuries at the injury site in the area of the endplate. Bone changes were minimal adjacent to the injury site and not observed in the far field. * indicate significant difference between groups with p<0.05.

### EP microfracture increased SubP in spinal dorsal horn

SubP, a pain-related neurotransmitter produced from nociceptive neurons, was mainly localized in laminae I and II of the spinal cord dorsal horn (Figure 7A). The percentage area of SubP-ir (relative to dorsal horn) was significantly increased in rat spinal dorsal horns from EP injury groups (3.61%±0.82% and 3.88%±0.69% for EP+PBS and EP+TNF groups, respectively) compared to that of Sham (2.15%±0.65%) (Figure 7C).

**Figure 7:**
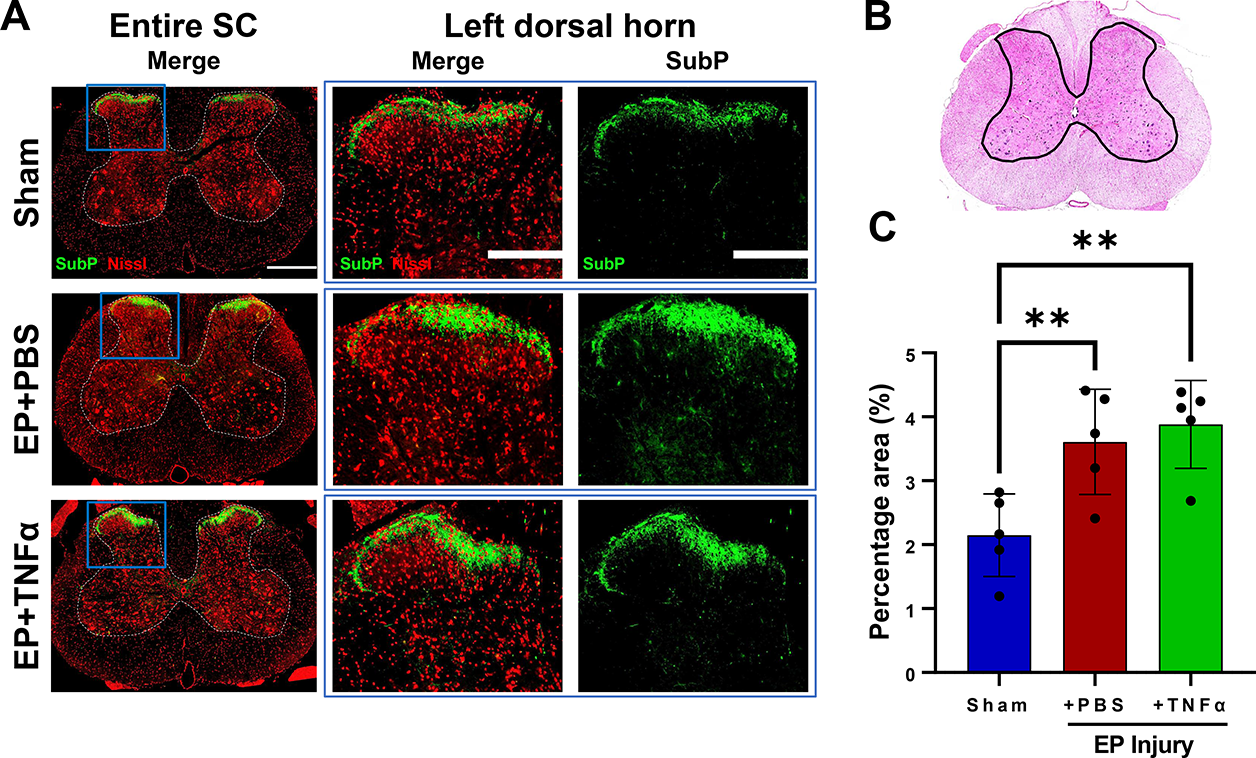
Spinal Cord sensitization from EP injury. A) Immunofluorescence images of spinal cord (SC) with outlined gray matter showing Neurons (red) and presence of substance P (green) in different groups with focus on dorsal horns. B) hematoxylin and eosin staining of spinal cord with outline of grey matter. C) quantification of SubP in dorsal horn. ** indicate significant difference between groups with p<0.01.

### NP T2 correlated with IVDD, SubP and pain-like behaviors

NP T2 relaxation times significantly correlated with histological IVD degeneration and SC SubP-ir indicating cross-talk between IVD and SC due to EP microfracture (Figure 8). NP T2 also correlated with hindpaw von Frey and axial grip force, indicating an association of NP T2 times with pain-like behaviors.

**Figure 8:**
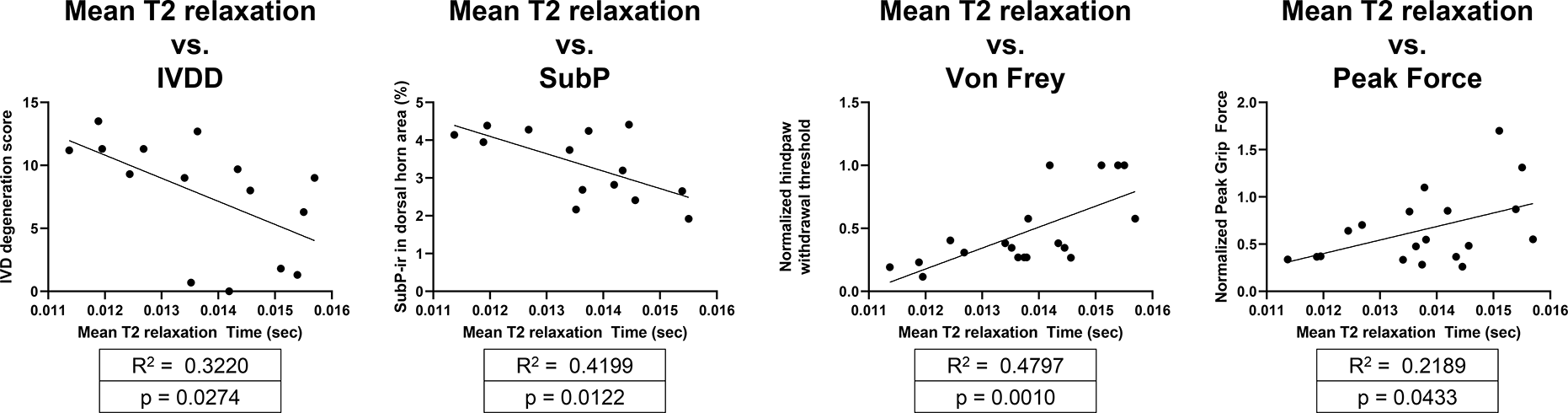
Correlations of NP T2 with IVDD, SC SubP, and pain-related behavioral measurements of von Frey and peak grip force.

## Discussion

Current clinical diagnoses for EP-driven IVD degeneration lacks phenotypic precision, has a high incidence of pain, and treatments have limited efficacy [50, 51]. A rat in vivo EP-driven IVD degeneration model was developed to better understand the progression of EP defects to EP-driven IVD degeneration, chronic pain and cross-talk between spinal tissues. We created a transcorporeal EP injury with a size of ~2% of the average rat lumbar EP surface area [52], which we describe as a microfracture injury. This study showed EP microfracture induced IVD degeneration and height loss, Modic-like changes, and increased pain-like behaviors with spinal cord sensitization. Pain-related behaviors and spinal cord SubP-ir significantly correlated with NP T2 relaxation time within the NP, suggesting EP microfracture injury resulted in pain associated with crosstalk between vertebrae, IVD, and spinal cord. The chronic and persistent pain-related behavior phenotype with hindpaw sensitivity and increased spinal cord SubP-ir revealed central sensitization, which suggests this vertebral injury with EP puncture induces broad changes to the entire spinal column that must all be considered during diagnosis and treatment.

This EP injury model was characterized using imaging modalities and defined MC presence and severity using MRI [46] that provides parallels to the human clinical condition. MCs are vertebral endplate and adjacent bone marrow lesions visible via MRI, including three phenotypes (MCs type1, MCs type2, MCs type3), first described by de Roos et al. [53] and Modic et al. [10]. MC type 1 fibrotic lesions have the highest association with pain, while MC type 3 sclerotic features are often asymptomatic [29]. EP injury is described to occur through traumatic fractures or accumulating microfracture and MCs. MCs prevalence is high in patients with back pain and EP injury [54–56]. MCs type1 reflect a state of active degeneration, and biomechanical instability of the lumbar spine and are considered a marker of active back pain with poor surgical prognosis [17, 29, 46, 57–59].MCs type 1 reflect inflammatory and fibrotic subchondral lesions with hypointense signal in T1w and hyperintense signal in T2w MR images.

In the current study, MC type 1-like changes were more easily visible when comparing EP+TNFα with Sham on T2w and T1w images than with EP+PBS group which were predominantly visible on T2w MRI. Nevertheless, MCs were apparent on MRI in all EP injured samples. Histology further demonstrated EP microfracture injury caused MC type 1-like changes with adjacent granulation tissue and inflammatory cell infiltration in the bone marrow that was more severe for EP+TNFα than EP+PBS (Figure 4; Supplementary Figure 2). The findings demonstrate TNFα injection might stimulate MC type 1-like changes, which is consistent with the results from Dudli et al indicating that proinflammatory stimulus was critical to induce MC1-like changes [29]. However, EP microfracture injury alone created crosstalk between vertebral bone marrow and IVD with inflammatory cell infiltration, suggesting that proinflammatory conditions were present in both EP injury groups. EP fractures can cause MCs with inflammatory and catabolic changes to the IVD, which are suggested to occur from an autoimmune response of the bone marrow against the IVD [18, 29, 31, 54, 55].

The pain-related behavioral phenotype was characterized with reduced axial grip strength and increased mechanical sensitivity at the hindpaw suggesting local and central pain [60]. PBS and TNFα injections had identical behavioral results, indicating that the behavioral phenotypes were affected by the presence of the EP puncture injury, and not the type of injectate. Results therefore suggest that the EP puncture injury itself causes pain or disability that may be a result of EP-injury induced axial instability or inflammation. The grip force assay requires contraction of axial musculature for the animal to stabilize and grip the bar. The significant reduction in grip force is most likely related to axial mechanical discomfort and/or disability since previous EP injury showed axial biomechanical instability in rat spinal segments in ex vivo biomechanical tests [61]. Results therefore suggest MC changes, as induced in this model, have an axial pain phenotype.

Central sensitization was demonstrated by enhanced mechanical hindpaw sensitivity, since there was no evidence that this vertebral EP puncture injury resulted in spinal cord or nerve root compression from herniation or vertebral remodeling on imaging or histology. However, inflammatory conditions were observed with EP injury, with the presence of inflammatory cells, and MC1-like changes with increased spinal cord SubP-ir 8 weeks after EP microfracture injury. The SubP in the spinal cord were mainly at laminae I and II which include terminations of nociceptive A-delta and C nerve fibers [62]. Spinal cord SubP is increased from direct spinal cord injury [63] or spinal transection [64], from diabetic neuropathy models [65], and also following paw inflammatory injury [66], suggesting spinal cord sensitization can occur from inflammatory and neuropathic sources in the spinal cord or periphery. The current study adds EP microfracture injury to the list of conditions resulting in spinal cord sensitization, and shows that EP microfracture injury can create a discogenic pain condition with crosstalk between vertebrae, IVD and spinal cord.

EP+PBS and EP+TNFα groups had similar pain-like behavioral responses even though EP+TNFα had more severe Modic-like changes and IVD degeneration score. Results therefore suggest that pain and disability is driven more by the presence of the EP injury rather than the severity of the injury, although it remains likely that the more severe EP+TNFα condition would reduce healing potential and perhaps cause greater dysfunction at longer time points in this model. In context of the literature, EP microfracture injury causes inflammatory and marrow changes and central spinal cord sensitization. The pain-related behavior in our model is therefore most likely related to biomechanical instability as well as inflammation, suggesting important parallels with the human clinical condition.

Imaging results allow us to confirm that the EP defect was a local microfracture injury that resulted in a broader inflammatory response. However, quantitative analyses of vertebral BV/TV or marrow T2 relaxation times did not detect statistical differences due to the limited sample size, particularly for μCT which occurred on a subset of the samples (3 sham; 3 EP+PBS and 5 EP+TNFα) because of a technical error when a batch of samples were mistakenly embedded for histology prior to scanning. Trabecular microstructure remodeling occurred following EP microfracture as apparent on imaging, and there was a suggestion that trabecular remodeling occurred throughout the entire affected vertebra, even though quantitative results were inconclusive. This study therefore has similarities to the human condition, as Senck et al., used μCT to visualize EP-driven IVD degeneration and demonstrated a local increase of trabecular thickness inferior to the EP collapse, and trabecular microstructure in the immediate vicinity of the collapse seems to be less organized, showing thicker trabeculae with a decreased trabecular length [22]. It is expected that BV/TV would be decreased in both EP injury groups due to bone absorption and remodeling caused by bone marrow/NP autoimmunity reaction.

Some limitations are important to highlight. This is a model system where EP injury was induced to create an EP microfracture with injury precision enhanced using one experienced spine surgeon and fluoroscopy guidance. There are similarities with clinical studies that include trauma-induced EP injuries and neuroinnervation occurring in peripheral and central areas that can contribute to back pain [12]. This study used male rats because of their larger size that made surgery slightly easier, and their more well-characterized von Frey mechanical sensitivity to IVD disruptions[24, 60]. Future studies are required to include female rats to improve the generalizability of these findings. Lastly, this is a descriptive model characterization study, and we hope this model will be able to gain further insights into discogenic pain with future blocking and therapeutic screening studies.

## Conclusions

This study established a rat in vivo model of EP microfracture that caused MC type 1-like changes, IVD degeneration, and spinal cord sensitization. The behavioral, radiological and histological phenotypes of this model were characterized with several similarities with the human clinical condition. The presence of the EP microfracture injury caused crosstalk between vertebrae, IVD, and spinal cord sensitization, with TNFα injection increasing injury severity. The pain-like behaviors indicated spinal pathology and central sensitization occurred suggesting axial biomechanical instability and inflammation resulted in pain, and these changes were more dependent on the presence of EP injury than injury severity. This study motivates assessments of sex-dependent changes, and the use of this model for blocking and therapeutic screening studies that will enable improved understanding of the cross-talk between vertebrae, IVD and spinal cord.

## Acknowledgement

This work was funded by grants from the National Institute of Arthritis and Musculoskeletal and Skin Diseases NIH/NIAMS grants R01AR078857 & R01AR080096. Development of the remmiRARE pulse sequence was funded by NIH/NIBIB grant R01EB019980.

## Figure Legends

**Graphical abstract**: Endplate (EP) injury induced IVD degeneration, Modic-like changes & mechanical allodynia and central sensitization.

---

**Graphical abstract.**
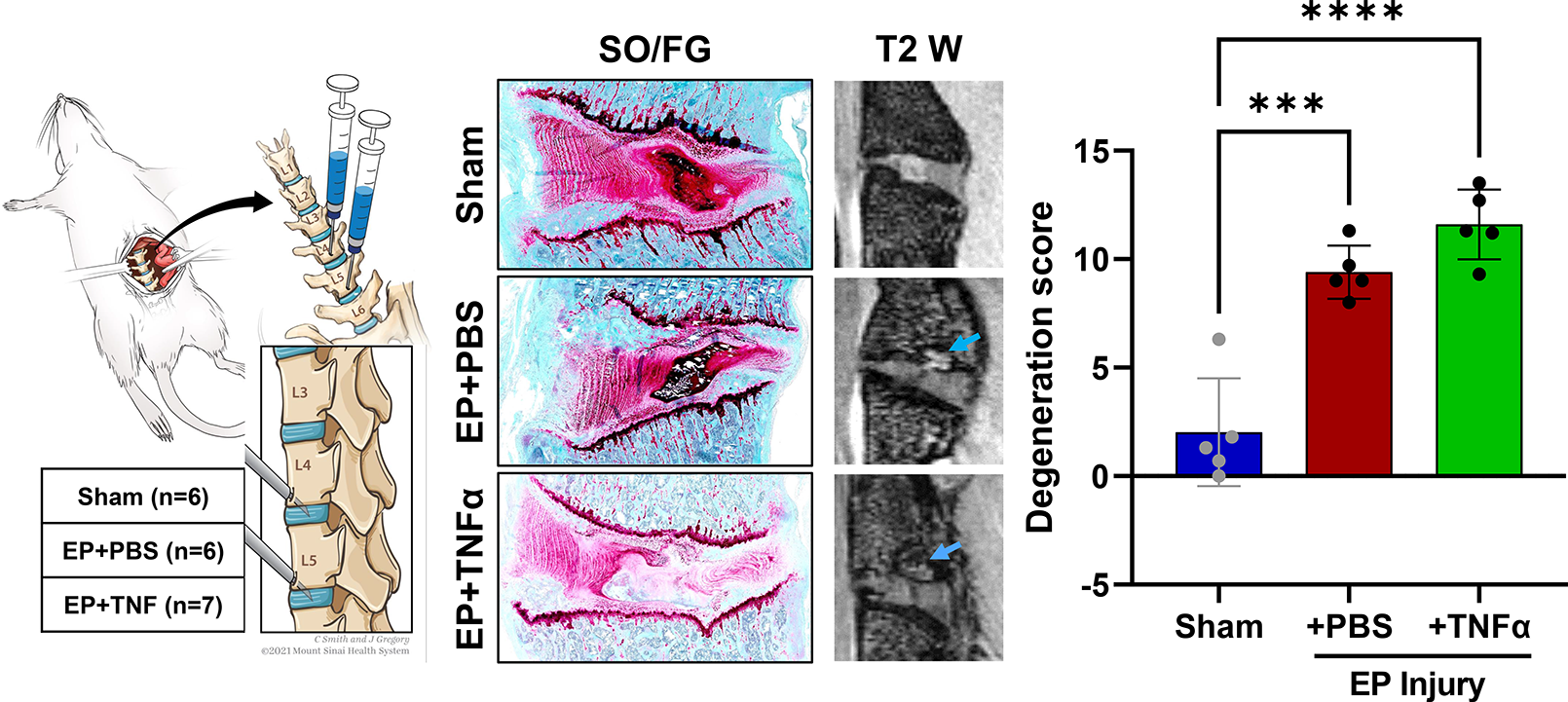

**Supplementary Figure 1:**
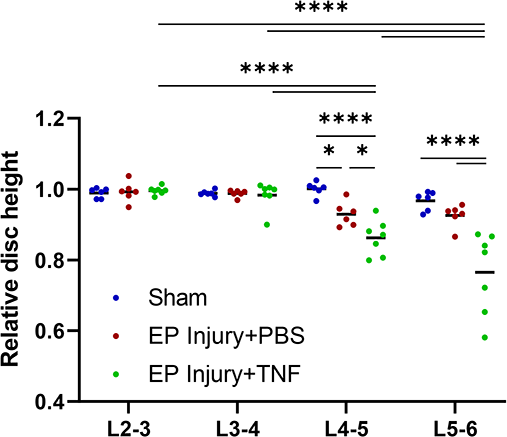
IVD height measured from Faxitron images showing effects of level. L2-3 and L3-4 were uninjured while L4-5 and L5-6 were injred levels. * and **** indicate significant differences with p<0.05 and p<0.0001 respectively.

**Supplementary Figure 2:**
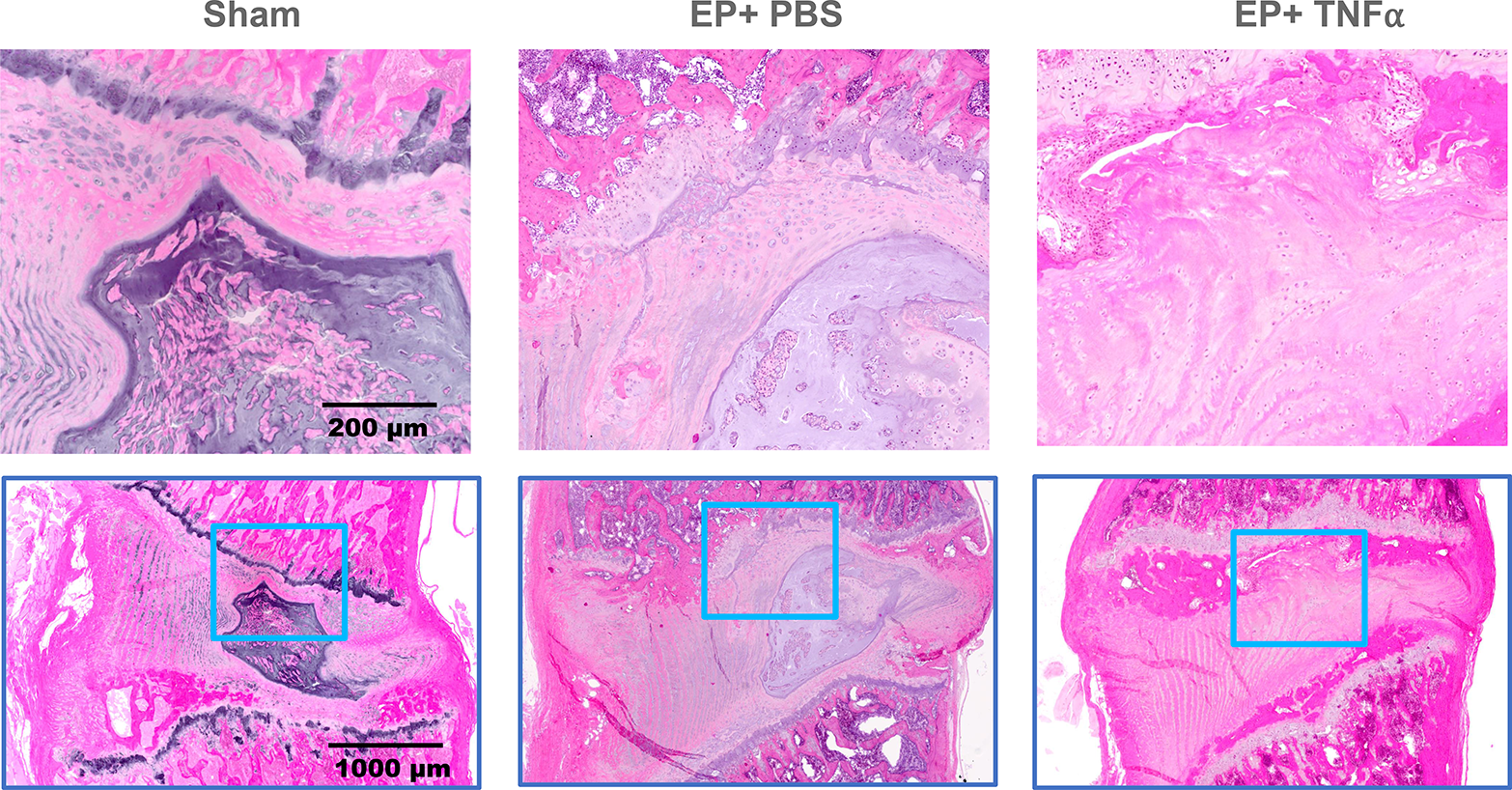
Hematoxylin and Eosin staining of IVDs showing disruption of IVD structure and inflammatory cell invasion into the IVD in both EP+PBS and EP+TNF injury groups.

## References

1. Adams MA, Dolan P. Intervertebral disc degeneration: evidence for two distinct phenotypes. J Anat. 2012;221(6):497–506.

2. Battie MC, Videman T, Levalahti E, Gill K, Kaprio J. Genetic and environmental effects on disc degeneration by phenotype and spinal level: a multivariate twin study. Spine (Phila Pa 1976). 2008;33(25):2801–8.

3. Hoy D, March L, Brooks P, et al. The global burden of low back pain: estimates from the Global Burden of Disease 2010 study. Ann Rheum Dis. 2014;73(6):968–74.

4. Wang D, Yuan H, Liu A, et al. Analysis of the relationship between the facet fluid sign and lumbar spine motion of degenerative spondylolytic segment using Kinematic MRI. Eur J Radiol. 2017;94:6–12.

5. Zehra U, Cheung JPY, Bow C, Lu W, Samartzis D. Multidimensional vertebral endplate defects are associated with disc degeneration, modic changes, facet joint abnormalities, and pain. J Orthop Res. 2019;37(5):1080–9.

6. Mallow GM, Zepeda D, Kuzel TG, et al. ISSLS PRIZE in Clinical Science 2022: Epidemiology, risk factors and clinical impact of juvenile Modic changes in paediatric patients with low back pain. Eur Spine J. 2022;31(5):1069–79.

7. Jensen TS, Karppinen J, Sorensen JS, Niinimaki J, Leboeuf-Yde C. Vertebral endplate signal changes (Modic change): a systematic literature review of prevalence and association with non-specific low back pain. Eur Spine J. 2008;17(11):1407–22.

8. Braten LCH, Rolfsen MP, Espeland A, et al. Efficacy of antibiotic treatment in patients with chronic low back pain and Modic changes (the AIM study): double blind, randomised, placebo controlled, multicentre trial. BMJ. 2019;367:l5654.

9. Herlin C, Kjaer P, Espeland A, et al. Modic changes-Their associations with low back pain and activity limitation: A systematic literature review and meta-analysis. PLoS One. 2018;13(8):e0200677.

10. Modic MT, Steinberg PM, Ross JS, Masaryk TJ, Carter JR. Degenerative disk disease: assessment of changes in vertebral body marrow with MR imaging. Radiology. 1988;166(1 Pt 1):193–9.

11. Yang X, Karis DSA, Vleggeert-Lankamp CLA. Association between Modic changes, disc degeneration, and neck pain in the cervical spine: a systematic review of literature. Spine J. 2020;20(5):754–64.

12. Lotz JC, Fields AJ, Liebenberg EC. The role of the vertebral end plate in low back pain. Global Spine J. 2013;3(3):153–64.

13. Maatta JH, Wadge S, MacGregor A, Karppinen J, Williams FM. ISSLS Prize Winner: Vertebral Endplate (Modic) Change is an Independent Risk Factor for Episodes of Severe and Disabling Low Back Pain. Spine (Phila Pa 1976). 2015;40(15):1187–93.

14. Albert HB, Kjaer P, Jensen TS, Sorensen JS, Bendix T, Manniche C. Modic changes, possible causes and relation to low back pain. Med Hypotheses. 2008;70(2):361–8.

15. Crock HV. Internal disc disruption. A challenge to disc prolapse fifty years on. Spine (Phila Pa 1976). 1986;11(6):650–3.

16. Dudli S, Boffa DB, Ferguson SJ, Haschtmann D. Leukocytes Enhance Inflammatory and Catabolic Degenerative Changes in the Intervertebral Disc After Endplate Fracture In Vitro Without Infiltrating the Disc. Spine (Phila Pa 1976). 2015;40(23):1799–806.

17. Dudli S, Fields AJ, Samartzis D, Karppinen J, Lotz JC. Pathobiology of Modic changes. Eur Spine J. 2016;25(11):3723–34.

18. Dudli S, Sing DC, Hu SS, et al. ISSLS PRIZE IN BASIC SCIENCE 2017: Intervertebral disc/bone marrow cross-talk with Modic changes. Eur Spine J. 2017;26(5):1362–73.

19. Kerttula LI, Serlo WS, Tervonen OA, Paakko EL, Vanharanta HV. Post-traumatic findings of the spine after earlier vertebral fracture in young patients: clinical and MRI study. Spine (Phila Pa 1976). 2000;25(9):1104–8.

20. Rajasekaran S, Babu JN, Arun R, Armstrong BR, Shetty AP, Murugan S. ISSLS prize winner: A study of diffusion in human lumbar discs: a serial magnetic resonance imaging study documenting the influence of the endplate on diffusion in normal and degenerate discs. Spine (Phila Pa 1976). 2004;29(23):2654–67.

21. Adams MA, Roughley PJ. What is intervertebral disc degeneration, and what causes it? Spine. 2006;31(18):2151–61.

22. Senck S, Trieb K, Kastner J, Hofstaetter SG, Lugmayr H, Windisch G. Visualization of intervertebral disc degeneration in a cadaveric human lumbar spine using microcomputed tomography. J Anat. 2020;236(2):243–51.

23. Krock E, Millecamps M, Currie JB, Stone LS, Haglund L. Low back pain and disc degeneration are decreased following chronic toll-like receptor 4 inhibition in a mouse model. Osteoarthritis Cartilage. 2018;26(9):1236–46.

24. Lai A, Moon A, Purmessur D, et al. Annular puncture with tumor necrosis factor-alpha injection enhances painful behavior with disc degeneration in vivo. Spine J. 2016;16(3):420–31.

25. Lai A, Ho L, Evashwick-Rogler TW, et al. Dietary polyphenols as a safe and novel intervention for modulating pain associated with intervertebral disc degeneration in an in-vivo rat model. PLoS One. 2019;14(10):e0223435.

26. Leimer EM, Gayoso MG, Jing L, Tang SY, Gupta MC, Setton LA. Behavioral Compensations and Neuronal Remodeling in a Rodent Model of Chronic Intervertebral Disc Degeneration. Sci Rep. 2019;9(1):3759.

27. Miyagi M, Ishikawa T, Orita S, et al. Disk injury in rats produces persistent increases in pain-related neuropeptides in dorsal root ganglia and spinal cord glia but only transient increases in inflammatory mediators: pathomechanism of chronic diskogenic low back pain. Spine (Phila Pa 1976). 2011;36(26):2260–6.

28. Mosley GE, Wang M, Nasser P, et al. Males and females exhibit distinct relationships between intervertebral disc degeneration and pain in a rat model. Sci Rep. 2020;10(1):15120.

29. Dudli S, Liebenberg E, Magnitsky S, Lu B, Lauricella M, Lotz JC. Modic type 1 change is an autoimmune response that requires a proinflammatory milieu provided by the ‘Modic disc’. Spine J. 2018;18(5):831–44.

30. Han C, Wang T, Jiang HQ, et al. An Animal Model of Modic Changes by Embedding Autogenous Nucleus Pulposus inside Subchondral Bone of Lumbar Vertebrae. Sci Rep. 2016;6:35102.

31. Cinotti G, Della Rocca C, Romeo S, Vittur F, Toffanin R, Trasimeni G. Degenerative changes of porcine intervertebral disc induced by vertebral endplate injuries. Spine (Phila Pa 1976). 2005;30(2):174–80.

32. Holm S, Holm AK, Ekstrom L, Karladani A, Hansson T. Experimental disc degeneration due to endplate injury. J Spinal Disord Tech. 2004;17(1):64–71.

33. Vadala G, Russo F, De Strobel F, et al. Novel stepwise model of intervertebral disc degeneration with intact annulus fibrosus to test regeneration strategies. J Orthop Res. 2018;36(9):2460–8.

34. Elliott DM, Sarver JJ. Young investigator award winner: validation of the mouse and rat disc as mechanical models of the human lumbar disc. Spine (Phila Pa 1976). 2004;29(7):713–22.

35. Hwang PY, Allen KD, Shamji MF, et al. Changes in midbrain pain receptor expression, gait and behavioral sensitivity in a rat model of radiculopathy. Open Orthop J. 2012;6:383–91.

36. Kim JS, Kroin JS, Buvanendran A, et al. Characterization of a new animal model for evaluation and treatment of back pain due to lumbar facet joint osteoarthritis. Arthritis Rheum. 2011;63(10):2966–73.

37. Kim JS, Kroin JS, Li X, et al. The rat intervertebral disk degeneration pain model: relationships between biological and structural alterations and pain. Arthritis Res Ther. 2011;13(5):R165.

38. Lai A, Moon A, Purmessur D, et al. Assessment of functional and behavioral changes sensitive to painful disc degeneration. J Orthop Res. 2015;33(5):755–64.

39. Miyagi M, Ishikawa T, Kamoda H, et al. Assessment of pain behavior in a rat model of intervertebral disc injury using the CatWalk gait analysis system. Spine (Phila Pa 1976). 2013;38(17):1459–65.

40. Mosley GE, Hoy RC, Nasser P, et al. Sex Differences in Rat Intervertebral Disc Structure and Function Following Annular Puncture Injury. Spine (Phila Pa 1976). 2019.

41. Rousseau MA, Ulrich JA, Bass EC, Rodriguez AG, Liu JJ, Lotz JC. Stab incision for inducing intervertebral disc degeneration in the rat. Spine. 2007;32(1):17–24.

42. Zhang KB, Zheng ZM, Liu H, Liu XG. The effects of punctured nucleus pulposus on lumbar radicular pain in rats: a behavioral and immunohistochemical study. J Neurosurg Spine. 2009;11(4):492–500.

43. Ponnappan RK, Markova DZ, Antonio PJ, et al. An organ culture system to model early degenerative changes of the intervertebral disc. Arthritis Res Ther. 2011;13(5):R171.

44. Evashwick-Rogler TW, Lai A, Watanabe H, et al. Inhibiting tumor necrosis factor-alpha at time of induced intervertebral disc injury limits long-term pain and degeneration in a rat model. JOR Spine. 2018;1(2).

45. Miyagi M, Millecamps M, Danco AT, Ohtori S, Takahashi K, Stone LS. ISSLS Prize winner: Increased innervation and sensory nervous system plasticity in a mouse model of low back pain due to intervertebral disc degeneration. Spine (Phila Pa 1976). 2014;39(17):1345–54.

46. Udby PM, Samartzis D, Carreon LY, Andersen MO, Karppinen J, Modic M. A definition and clinical grading of Modic changes. J Orthop Res. 2022;40(2):301–7.

47. Smith SM, Jenkinson M, Woolrich MW, et al. Advances in functional and structural MR image analysis and implementation as FSL. Neuroimage. 2004;23 Suppl 1:S208–19.

48. Jenkinson M, Beckmann CF, Behrens TE, Woolrich MW, Smith SM. Fsl. Neuroimage. 2012;62(2):782–90.

49. Lai A, Gansau J, Gullbrand SE, et al. Development of a standardized histopathology scoring system for intervertebral disc degeneration in rat models: An initiative of the ORS spine section. JOR Spine. 2021;4(2):e1150.

50. Hao L, Li S, Liu J, Shan Z, Fan S, Zhao F. Recurrent disc herniation following percutaneous endoscopic lumbar discectomy preferentially occurs when Modic changes are present. J Orthop Surg Res. 2020;15(1):176.

51. Lurie JD, Moses RA, Tosteson AN, et al. Magnetic resonance imaging predictors of surgical outcome in patients with lumbar intervertebral disc herniation. Spine (Phila Pa 1976). 2013;38(14):1216–25.

52. Jaumard NV, Leung J, Gokhale AJ, Guarino BB, Welch WC, Winkelstein BA. Relevant Anatomic and Morphological Measurements of the Rat Spine: Considerations for Rodent Models of Human Spine Trauma. Spine (Phila Pa 1976). 2015;40(20):E1084–92.

53. de Roos A, Kressel H, Spritzer C, Dalinka M. MR imaging of marrow changes adjacent to end plates in degenerative lumbar disk disease. AJR Am J Roentgenol. 1987;149(3):531–4.

54. Adams MA, Freeman BJ, Morrison HP, Nelson IW, Dolan P. Mechanical initiation of intervertebral disc degeneration. Spine (Phila Pa 1976). 2000;25(13):1625–36.

55. Joe E, Lee JW, Park KW, et al. Herniation of cartilaginous endplates in the lumbar spine: MRI findings. AJR Am J Roentgenol. 2015;204(5):1075–81.

56. Theodorou DJ, Theodorou SJ, Kakitsubata S, Nabeshima K, Kakitsubata Y. Abnormal Conditions of the Diskovertebral Segment: MRI With Anatomic-Pathologic Correlation. AJR Am J Roentgenol. 2020;214(4):853–61.

57. Kaapa E, Luoma K, Pitkaniemi J, Kerttula L, Gronblad M. Correlation of size and type of modic types 1 and 2 lesions with clinical symptoms: a descriptive study in a subgroup of patients with chronic low back pain on the basis of a university hospital patient sample. Spine (Phila Pa 1976). 2012;37(2):134–9.

58. Rahme R, Moussa R. The modic vertebral endplate and marrow changes: pathologic significance and relation to low back pain and segmental instability of the lumbar spine. AJNR Am J Neuroradiol. 2008;29(5):838–42.

59. Splendiani A, Bruno F, Marsecano C, et al. Modic I changes size increase from supine to standing MRI correlates with increase in pain intensity in standing position: uncovering the “biomechanical stress” and “active discopathy” theories in low back pain. Eur Spine J. 2019;28(5):983–92.

60. Mosley GE, Evashwick-Rogler TW, Lai A, Iatridis JC. Looking beyond the intervertebral disc: the need for behavioral assays in models of discogenic pain. Ann N Y Acad Sci. 2017;1409(1):51–66.

61. Wang D, Lai A, Gansau J, et al. Ex vivo biomechanical evaluation of Acute lumbar endplate injury and comparison to annulus fibrosus injury in a rat model. J Mech Behav Biomed Mater. 2022;131:105234.

62. Merighi A, Carmignoto G, Gobbo S, et al. Neurotrophins in spinal cord nociceptive pathways. Prog Brain Res. 2004;146:291–321.

63. Kim JS, Ahmadinia K, Li X, et al. Development of an Experimental Animal Model for Lower Back Pain by Percutaneous Injury-Induced Lumbar Facet Joint Osteoarthritis. J Cell Physiol. 2015;230(11):2837–47.

64. Zachariou V, Goldstein BD. Dynorphin-(1-8) inhibits the release of substance P-like immunoreactivity in the spinal cord of rats following a noxious mechanical stimulus. Eur J Pharmacol. 1997;323(2–3):159–65.

65. Wan FP, Bai Y, Kou ZZ, et al. Endomorphin-2 Inhibition of Substance P Signaling within Lamina I of the Spinal Cord Is Impaired in Diabetic Neuropathic Pain Rats. Front Mol Neurosci. 2016;9:167.

66. Zucoloto AZ, Manchope MF, Borghi SM, et al. Probucol Ameliorates Complete Freund’s Adjuvant-Induced Hyperalgesia by Targeting Peripheral and Spinal Cord Inflammation. Inflammation. 2019;42(4):1474–90.

